# Linking neural population formatting to function

**DOI:** 10.1101/2025.01.03.631242

**Authors:** Douglas A. Ruff, Sol K. Markman, Jason Z. Kim, Marlene R. Cohen

## Abstract

Animals capable of complex behaviors tend to have more distinct brain areas than simpler organisms, and artificial networks that perform many tasks tend to self-organize into modules (*1–3*). This suggests that different brain areas serve distinct functions supporting complex behavior. However, a common observation is that essentially anything that an animal senses, knows, or does can be decoded from neural activity in any brain area (*4–6*). If everything is everywhere, why have distinct areas? Here we show that the function of a brain area is more related to how different types of information are combined (formatted) in neural representations than merely whether that information is present. We compared two brain areas: the middle temporal area (MT), which is important for visual motion perception (*7, 8*), and the dorsolateral prefrontal cortex (dlPFC), which is linked to decision-making and reward expectation (*9, 10*)). When monkeys based decisions on a combination of motion and reward information, both types of information were present in both areas. However, they were formatted differently: in MT, they were encoded separably, while in dlPFC, they were represented jointly in ways that reflected the monkeys’ decision-making. A recurrent neural network (RNN) model that mirrored the information formatting in MT and dlPFC predicted that manipulating activity in these areas would differently affect decision-making. Consistent with model predictions, electrically stimulating MT biased choices midway between the visual motion stimulus and the preferred direction of the stimulated units (*11*), while stimulating dlPFC produced ‘winner-take-all’ decisions that sometimes reflected the visual motion stimulus and sometimes reflected the preference of the stimulated units, but never in between. These results are consistent with the tantalizing possibility that a modular structure enables complex behavior by flexibly reformatting information to accomplish behavioral goals.

## The formatting of different information sources in neural population responses is not obvious from single neurons

It has long been known that the responses of individual neurons reflect multiple sensory, cognitive, and/or motor processes. For example, MT neurons are tuned for visual motion direction (*7, 8, 12–14*) and their responses are modulated (often multiplicatively scaled) by reward information (e.g. the expected reward associated with a stimulus or choice) and other cognitive processes (*15–18*).

However, known patterns of tuning and modulation gleaned from single neuron studies are consistent with multiple ways of formatting information about motion direction and reward information in the population (which is sometimes called representational geometry or neural population geometry (*19, 20*)). The different possibilities arise because even identically tuned neurons have heterogeneous modulation by cognitive processes. Simulating this heterogeneity by adding some randomness in the amount of modulation by reward expectation among neurons with identical tuning for motion direction (Fig. 1A; methods) can produce population representations of motion direction and reward expectation that are either separable (encoded in different dimensions in a space where the response of each neuron is one dimension; Fig. 1B, C, D) or combined (encoded in the same dimensions; Fig. 1E, F, G). The difference between separable and combined population formatting is not knowable from single neuron responses and instead arises from how and whether modulation by reward expectation is coordinated across the population.

**Figure 1.**
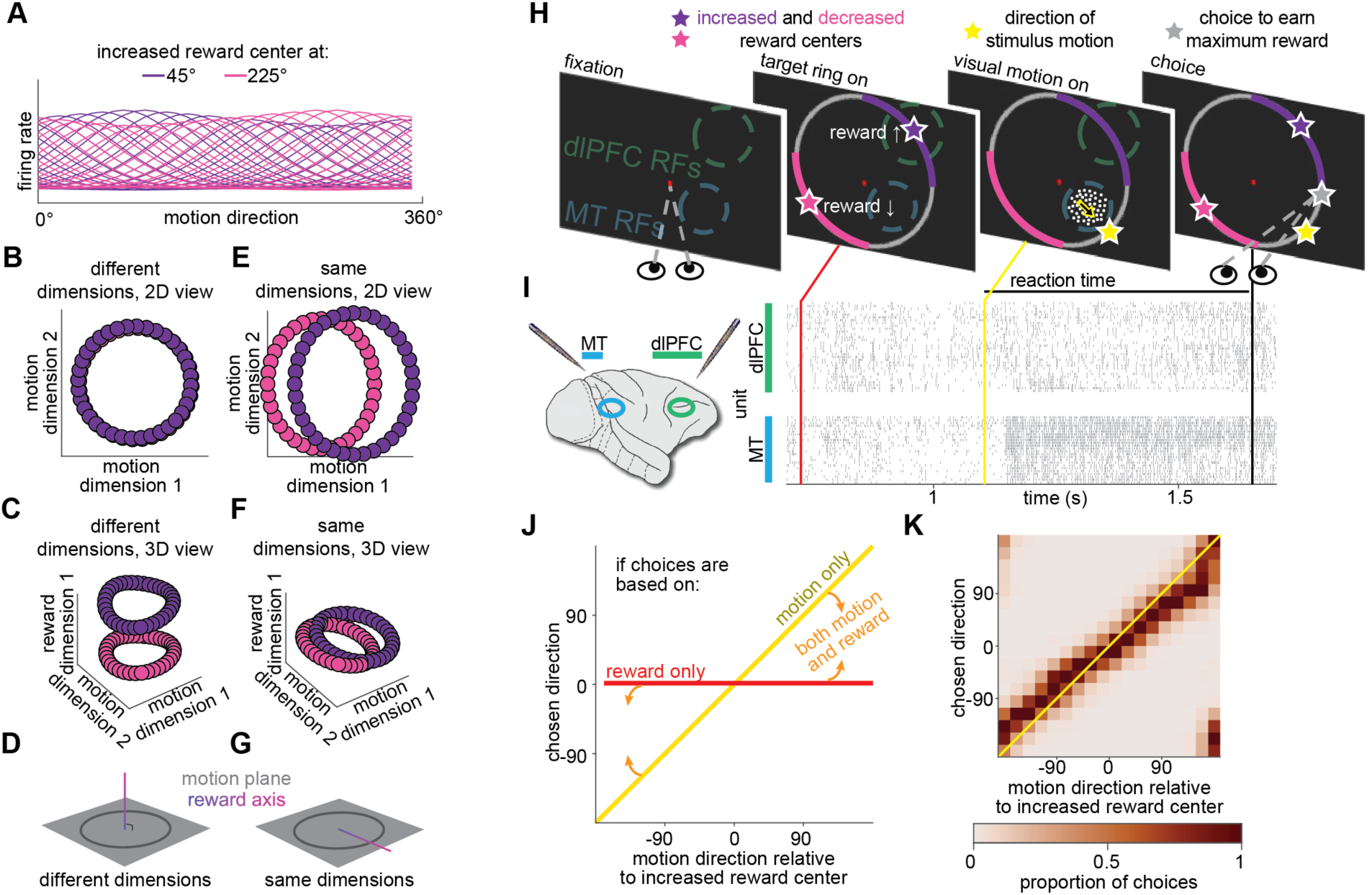
Population formatting, task and behavior. **A)** Simulation demonstrating that the formatting of different information in neural population responses cannot be inferred from single neurons. Subsets of tuning functions from simulated neurons whose preferred visual motion directions are uniformly distributed across 360° of motion in 10° steps and whose gain depends on the difference between their preferred motion direction and the direction of maximum expected reward during a behavioral task (here either 45° or 225°; details inspired from single neuron studies (31)). The modulation for each neuron was drawn from a distribution whose mean is shown in (A; Methods). **B)** Example population representation of motion direction and reward information from the simulated population in A (using one representative random draw from the distribution of reward modulations) where motion direction and reward information are encoded separably (so they are in different dimensions of the space where the response of each neuron is one dimension). For illustration, the population responses are projected into the two dimensions that encode the most visual motion direction information. **C)** Same as (B), with a third dimension that accounts for the most reward information. Colors denote the two reward conditions depicted in (A). **D)** Schematic showing population formatting where motion direction and reward expectation are represented separably (different, or orthogonal, dimensions). **E)** Example population representation of motion direction information and reward information from the simulated population in (A) using a different random draw from the same distribution of reward modulations as in (B), picked for illustration because this random draw produced information about motion direction and reward expectation that are combined (occur in the same dimensions of neural population space). Conventions as in (B). **F)** Same as (E), with a third dimension that accounts for the most reward information. Conventions as in (C). **G)** Schematic showing population formatting where reward expectation and motion direction are represented jointly (same dimensions). **H)** Continuous estimation task schematic. Monkeys indicated their choice by looking at a point on the target ring that corresponded to a direction judgment. They were rewarded according to the accuracy of their estimate of the direction of a dynamic random dot display (actual motion direction depicted by yellow star, gray star indicates choice that would result in maximum for this trial), which was scaled by a factor indicated by the color of that portion of the target ring (see Methods and Supplementary Figure 1; pink and purple colored stars signify the center of these regions, no stars were shown to the monkeys). The motion direction and reward condition were independently randomly interleaved from trial to trial, and the target ring colors associated with increased or decreased reward varied in blocks of trials with uncued changes. **I)** We recorded populations of neurons in MT and dlPFC using multielectrode probes. In the raster plot, each tick represents an action potential from an example trial, each row is a multiunit site in dlPFC (top) or MT (bottom). The x-axis represents time (target ring onset, red; visual motion stimulus onset, yellow; onset of eye movement indicating choice, black). **J)** Schematic depicting psychometric functions when choices are based on visual motion direction (yellow), reward condition (red), or a combination (orange arrows signifying choices that fall in between the yellow and red lines). **K)** Behavior from 27 experimental sessions from two monkeys (18 from monkey 1; 9 from monkey 2; 41,106 total choices) demonstrate that the monkeys based their choices on a combination of motion direction and reward information.

Previous studies have found that the formatting of information from different sensory, cognitive, or motor sources varies across brain areas, tasks, and species (*20–29*). Artificial networks that are trained to perform many tasks or complex tasks tend to self-organize into modules that likely format information differently (*1, 2*). The goal of our study was to understand the relationship between these different formatting schemes and the function of a neural population.

## New insights from a continuous, information-integration task

Natural decision-making provides inspiration about how to elicit rich enough behavior to reveal strong links between decisions and neural population responses. Natural behavior often requires decision-makers to make continuous estimates that reflect a combination of information from disparate sensory and cognitive sources. For example, one might estimate the ripeness of a peach by combining information about its color and prior knowledge about the quality of fruit available at a particular farm stand.

Continuous estimation tasks provide richer behavioral outputs when compared to typical binary choice laboratory tasks (see also (*11, 30*)), which enhances the statistical ability to test hypotheses about the relationship between neuronal responses, causal manipulations, and behavior.

We used a controlled laboratory task that mimics these aspects of natural decisions. Two adult rhesus monkeys (Macaca mulatta, both male, 8 and 10 kg) performed a continuous estimation task in which they combined sensory information (motion direction) with cognitive information (the reward magnitude associated with different choices). They earned different quantities of fruit juice reward on each trial, based on the accuracy of their motion direction estimate and a scaling factor associated with particular choices. Colored portions of the target ring the monkeys used to indicate their choice determined which choices would be associated with increased, decreased, or unchanged scaling of the reward earned from the accuracy of their visual motion estimate (Figure 1H, I, and Supplementary Figure 1). We refer to the center of the colored regions as the increased reward or decreased reward center and different locations of those regions (which varied from trial to trial) as reward conditions. The monkeys demonstrated that they integrated motion and reward information by making choices that varied smoothly with the direction of the visual stimulus and biasing choices toward larger expected rewards (Figure 1J, K; larger rewards were typically accompanied by faster reaction times; Supplementary Figure 2).

## Motion and reward information are formatted differently in visual and frontal cortex

While the monkeys performed the continuous estimation task, we recorded simultaneously from a few dozen neurons in two brain areas thought to contribute differently to the decision computation (Figure 1I). Neurons in area MT are critical for motion perception (*7, 8, 11*) and neurons in dlPFC are part of a network involved in decision-making, reward processing, and eye movement planning (*9, 10, 32*). Consistent with observations that sensory, cognitive, and motor information is encoded throughout the brain (*5, 6, 33, 34*), motion and reward information were readily apparent in the population activity of both MT and dlPFC (Figure 2) and dlPFC units tended to exhibit more non-linear mixed selectivity for motion and reward than MT units ((*19, 35*); Supplementary Figure 3).

**Figure 2:**
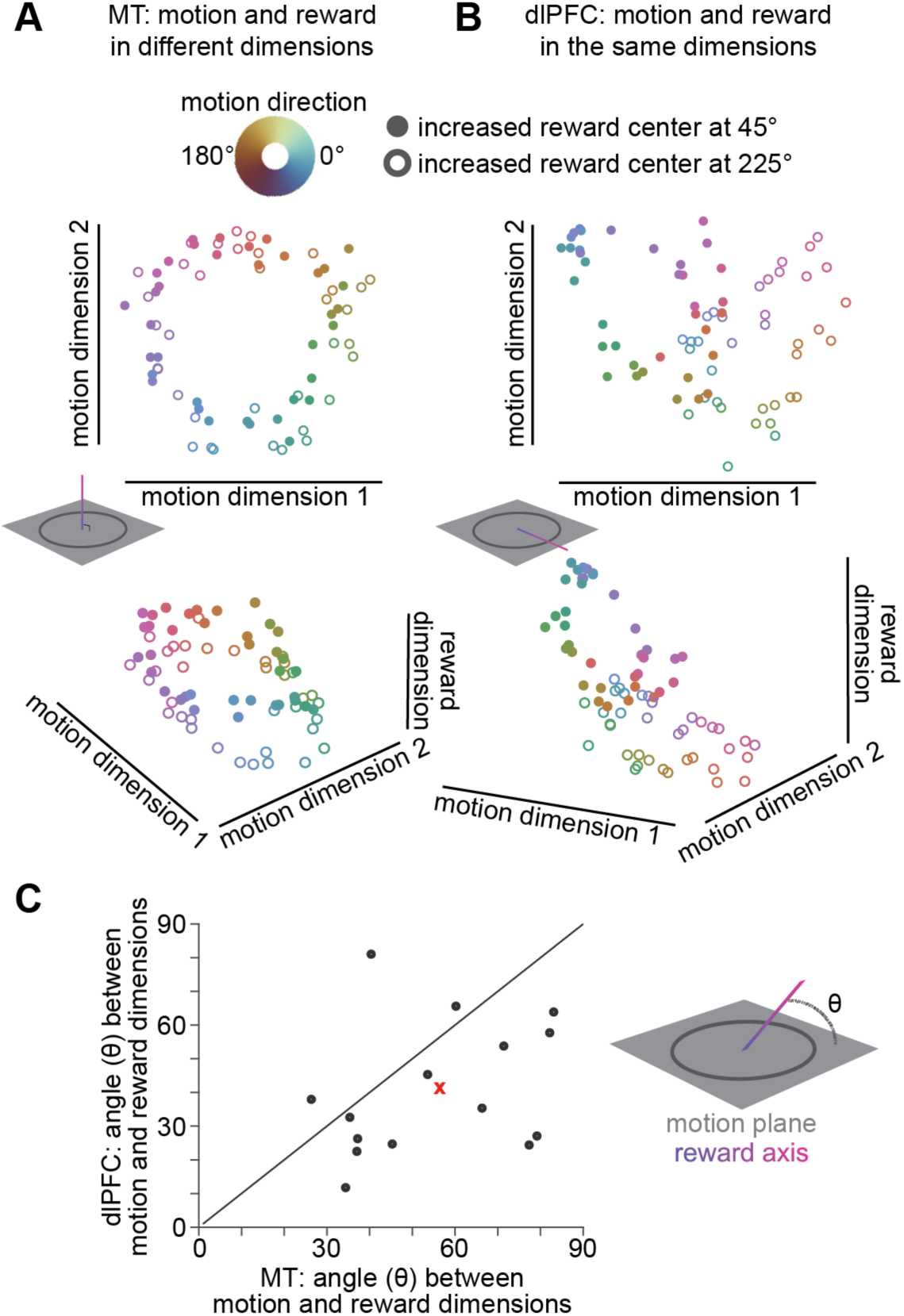
Both MT and dlPFC encode motion and reward information, but format that information differently. **A)** MT population responses from an example recording session projected onto the two axes of neural population space that account for the most variance in motion direction (top) and also the most variance across reward conditions (bottom; z-axis). Each point is the mean population response to one motion direction (colors), and open and filled circles are from trials in different reward conditions. **B)** Same as (A), for dlPFC population responses. **C)** Motion and reward information are more separable in MT than dlPFC. In 15 sessions in which MT and dlPFC neurons were recorded simultaneously, the angle between the dimensions accounting for the most variance in motion direction and reward condition (𝜃 in the schematic) were larger in MT (x-axis) than in dlPFC (y-axis; Wilcoxon signed-rank test, p<0.05). The red x represents the mean across all 15 sessions.

However, motion and reward information were formatted differently in the two areas. In MT, population representations of motion and reward tended to be separable, such that information about them occupied distinct dimensions of a neural population space where each dimension represents the response of one neuron (Figure 2A). In contrast, in dlPFC, motion and reward tended to be represented jointly, such that they occupied overlapping dimensions in neural population space (Figure 2B). This difference in neural population formatting (Figure 2C) means that motion and reward information can each be decoded independently from MT, while in dlPFC, the same population activity can indicate different motion directions in different reward conditions. These differences in formatting make this a good system in which to investigate questions about how formatting is related to the contribution of a brain area to behavior.

## An artificial neural network to generate causal hypotheses

We sought to use causal manipulations (e.g. measuring the impact of perturbing neural activity on behavior) to understand the functional implications of differently formatted population representations. However, from the existing literature, it was difficult to generate hypotheses about how the outcomes of causal experiments would relate to information formatting. Some previous studies have attempted to link neural population formatting to information coding or theoretical ideas about how that information could, in principle, be read out (*36–38*). However, to our knowledge, no studies have linked neural population formatting directly to behavior by comparing, in either model or experiment, the effect of causally manipulating brain areas with different formatting.

To generate hypotheses that we could test in further causal experiments, we constructed a recurrent neural network (RNN) that reproduced the neural population formatting in MT and dlPFC (Figure 3A). This model took as inputs the same information available to the monkey (motion and reward information). We first trained an MT-like module to receive motion direction and the locations of increased and decreased reward centers as inputs and to output the motion direction and reward center locations, independently.

**Figure 3:**
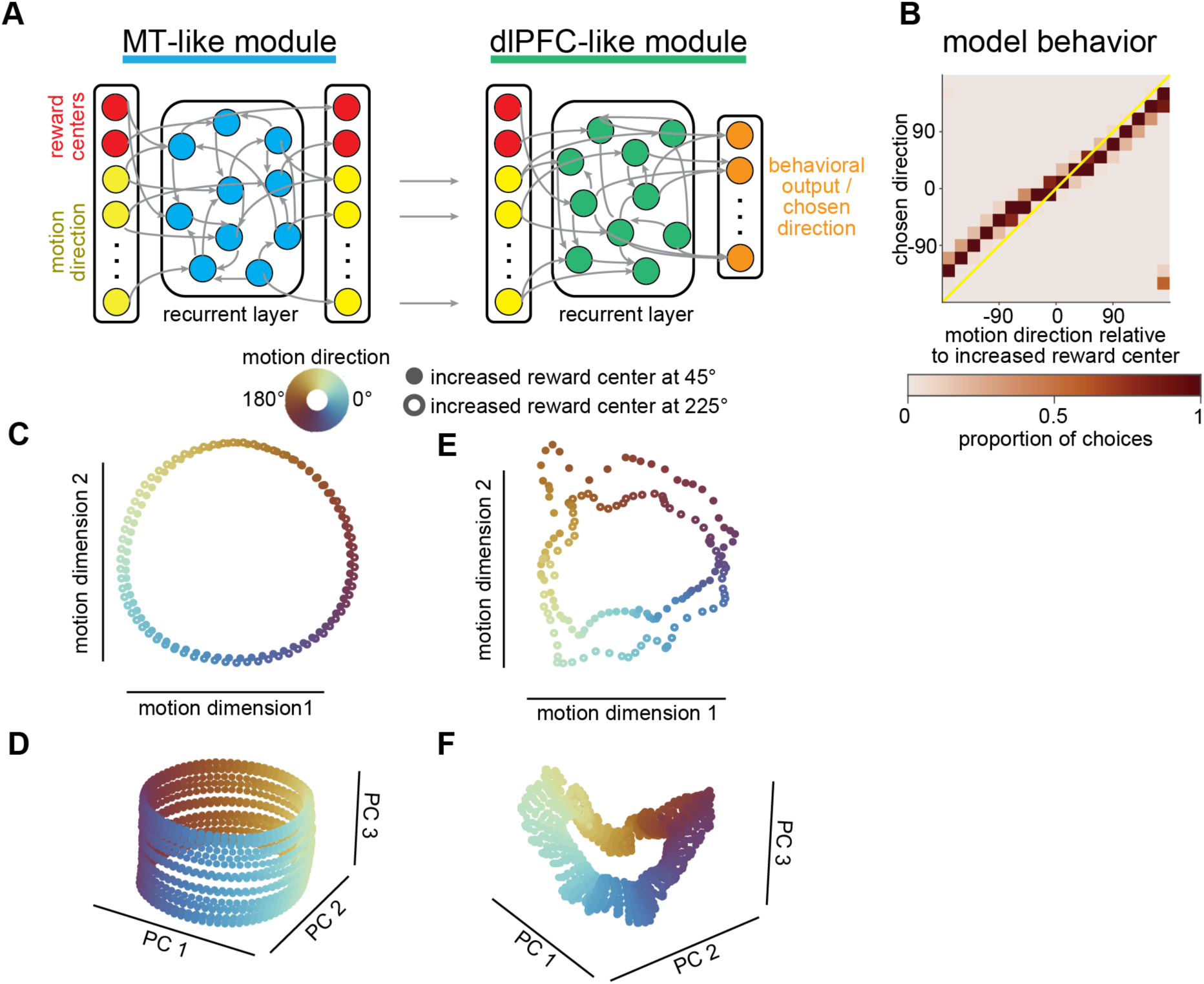
A two-stage artificial neural network replicates monkey behavior and population representations in MT and dlPFC. **A)** We constructed an RNN that reproduced neural population formatting in MT and dlPFC and performed the same task as the monkeys. An MT-like module was trained to output the direction of a noisy motion stimulus (yellow) and the increased reward center from two possible locations (red), in separable dimensions. The dlPFC-like module was trained to take the output of the MT-like module and reward condition on each trial and output an estimate of the reward associated with each possible choice. We take the model’s choice to be the direction with the largest predicted reward according to the output units of the dlPFC-like module (orange). **B)** Like the monkeys, the model makes choices that reflect both motion direction and reward information (compare to Figure 1J, K). Conventions as Figure 1K. **C)** Fixed points from the MT-projected onto the first two axes of population space that account for the most variance in motion direction (compare to Figure 2A). Colors represent different input motion directions and open and filled circles represent each of two possible reward conditions (increased reward center at either 45° or 225°). **D)** Same as C but including additional reward center positions projected onto the first three principal components (compare to Figure 2A). **E)** Same as C, for the dlPFC-like module. **F)** Same as D, for the dlPFC-like module (compare to Figure 2B).

After training, we froze the recurrent connections and made the motion output units of the MT-like module, along with the reward center locations, the inputs to the dlPFC-like module. This module was trained to output an estimate of the reward associated with each possible choice given the motion direction and reward condition. We took the model’s choice to be the direction associated with the highest expected reward (Figure 3B).

The model has many similarities to the data. Like the monkeys, it appropriately combined information about motion and reward (qualitatively compare Figure 3B to Figure 1K). By construction, the recurrent layers in the two modules represent motion and reward in ways that mimic MT and dlPFC, respectively. The MT-like module represented motion and reward information separably (Figure 3C, D), while these features were integrated in the dlPFC-like module (Figure 3E, F). This can be visualized by viewing the stable fixed points which the activity of the hidden layer approaches at the end of a trial (one fixed point for each combination of motion direction and the two reward conditions, Figure 3C, E; and, to more fully visualize the population structure, across a larger range of possible reward conditions; Figure 3D, F ). The similarities between the model and the data make this model a good tool for generating hypotheses about the relationship between the population formatting and behavior.

## Perturbing units in the recurrent layers illuminates the different functional roles of the two modules

We used our models to generate causal hypotheses about how the impact of perturbing the activity of model units (*39*) on choices depends on the way their module formats information (see Methods).

Perturbing units in the MT-like and dlPFC-like modules had qualitatively different effects on model choices. Consistent with results from a previous study that electrically stimulated MT (*11*), activating the MT-like module biased choices toward the preferred direction of the activated unit, provided that the difference between the motion direction and the preference of the activated unit was less than approximately 150° (vector averaging; Figure 4A-C, column 1). Perturbing the dlPFC-like module was very different. On some ostensibly identical trials, the model chose the motion direction and on others the preferred direction of the activated unit, and essentially never in between (winner take all; Figure 4A-C, column 3). Further, the behavioral impact of perturbation depended on the reward condition in the dlPFC-like module but not the MT-like module (compare Figure 4B and Figure 4C).

**Figure 4:**
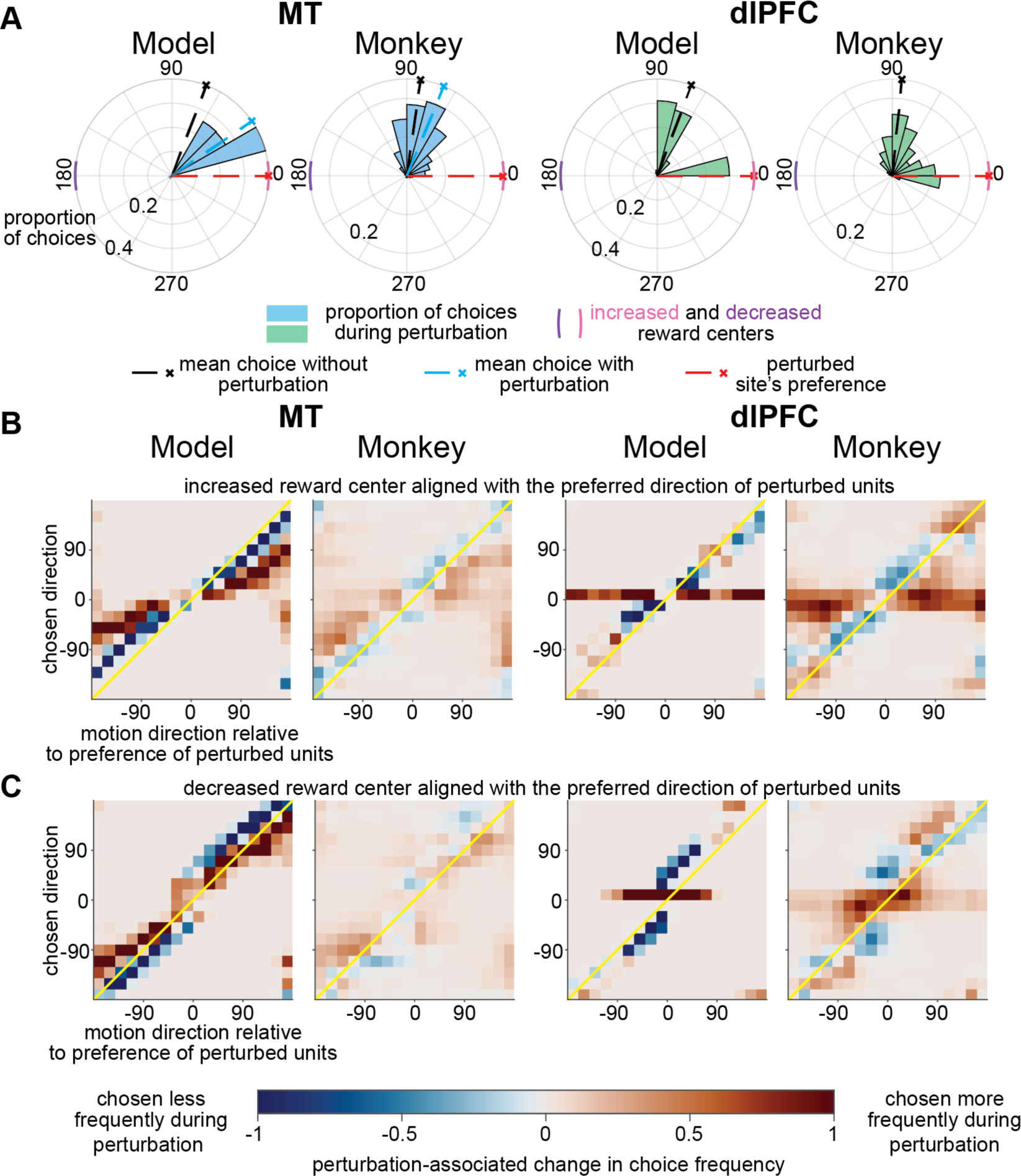
Consistent with model predictions, manipulating MT activity has qualitatively different effects on choices than manipulating dlPFC activity. **A)** Polar histograms of choices from model and monkey in area MT (columns 1 and 2) and dlPFC (columns 3 and 4) on an illustrative subset of trials in which the visual motion direction was 100° counterclockwise relative to the preferred direction of the perturbed units. The pink and purple arcs signify the increased and decreased reward centers, respectively. **B)** Perturbation-related difference in choices when the preference of the perturbed site was aligned with increased reward center for the model (columns 1 and 3) and experimental data (columns 2 and 4). The proportion of choices from during non-perturbation trials were subtracted from the proportion of choices during perturbation trials (Supplementary Figures 4 and 5 display choices from all conditions). In both the MT-like module and in monkey MT, perturbation biased choices toward the preference of the perturbed site (away from the identity line in columns 1 and 2; monkey data from 29 experimental sessions, comprising 50 sites; some sessions had stimulation on different electrodes on interleaved trials). In both the dlPFC-like module and in monkey dlPFC, perturbation produced bimodal choices that were directed either to the preference the perturbed site or to the mean choice during non-perturbation trials (resulting in choices clustered along the horizontal line representing the preference of the perturbed units in columns 3 and 4; monkey data from 23 experimental sessions, comprising 33 stimulated sites). **C)** Perturbation-related difference in choices when the preference of the perturbed site was aligned with the decreased reward center. Conventions as in (B).

## Manipulating MT and dlPFC activity had qualitatively different effects on choices

We tested the predictions of our model by electrically microstimulating one electrode at a time in MT or dlPFC on (or neither) randomly interleaved trials. Consistent with model predictions, microstimulating MT and dlPFC affected choices in distinct ways. As in previous work (*11*) and as predicted by our model, microstimulating MT biased choices midway between the visual stimulus motion direction and the preferred direction measured at the stimulated site (vector averaging; Figure 4A-C, column 2). In contrast, and consistent with the prediction of our model, microstimulating dlPFC produced a bimodal choice distribution, where the monkeys sometimes reported the direction of the visual motion and other times reported the preference measured at the stimulated site (winner take all; Figure 4A-C, column 4).

Furthermore, and consistent with the predictions of our model, stimulating MT and dlPFC differently interacted with the reward condition. The behavioral impact of MT microstimulation did not depend strongly on whether the increased or decreased reward was aligned with the preference of the cells recorded on the stimulating electrode (compare Figure 4B and 4C, column 2). The intuition is that because different information sources are encoded approximately independently in MT, there is no interaction between microstimulation and other information sources, like reward expectation. In contrast, in dlPFC, representations are joint such that the behavioral impact of microstimulation is much greater when the increased reward center is aligned with the preference of the stimulated site (compare Figure 4B and 4C, column 4).

Interestingly, the requirement of our task that monkeys combine information sources (from both visual motion and reward expectation) may have heightened the difference between MT and dlPFC stimulation. Unlike previous studies that elicited winner take all behavior using MT microstimulation when the angle between the visual motion and the preference of stimulated site was large (*11, 40*), we saw vector averaging whenever we stimulated MT in our main experiment. We hypothesize that the reward scaling encouraged the monkey to bias choices towards a larger reward even when the visual motion evidence suggested a choice associated with a smaller reward. Indeed, when we ran a subset of experiments with no reward scaling, we occasionally observed winner take all effects during MT microstimulation when the difference between the direction of the visual motion and the preference of the stimulated site was large (>150°).

Two controls highlight the robustness of the different impact of MT and dlPFC microstimulation on behavior. First, we controlled for the possibility that the impact of stimulation relative to the visual stimulus was larger in one brain area than the other, which led to qualitatively different impact on behavior. We controlled for this by changing the relative strength of the visual and electrical stimuli by reducing the visual motion coherence. Rather than qualitatively changing the impact of MT or dlPFC stimulation, this manipulation simply enhanced the impact of microstimulation in both areas, more strongly biasing choices towards the preference of the stimulated site in MT and increasing the proportion of trials on which the monkey reported the preference of the stimulated site in dlPFC. The qualitative difference between the two areas remained even at low motion coherence (Supplementary Figure 6 and 7).

Second, we controlled for the fact that for most experiments, the visual motion stimulus overlapped the MT receptive fields and the target ring overlapped the response fields of the dlPFC neurons. We controlled for this difference by conducting a subset of microstimulation experiments when the MT receptive fields overlapped the target ring rather than the visual motion stimulus. In these experiments, we did not systematically bias choices or induce winner-take-all behavior (Supplementary Figure 8).

## A causal link between neural population formatting and function

Our study made a leap from correlation to causation in the understanding the relationship between the formatting of neural population responses and behavior. As in previous studies (*40–44*), we found that when choices are based on both motion and reward information, both MT and dlPFC encode both information sources. But we demonstrated that those representations are formatted differently. This observation meant that this system is a platform for testing predictions about what neural population formatting implies about the function of those areas. A growing body of theoretical work has quantified how the flexibility or robustness in the way that information could, in principle, be read out to guide behavior depends on population formatting. (*45–48*). Indeed, this theoretical framework predicts that the formatting differences between MT and dlPFC are consistent with longstanding ideas that visual cortex functions as a close to veridical representation of the world and that dlPFC flexibly combines information to guide decision-making (*9, 10, 43*). However, we showed using our model trained only to reproduce the formatting that there is a causal link between formatting and function: as predicted by our models, electrically stimulating those areas had a qualitatively different impact on behavior. Our results support the exciting idea that the contributions of less well-studied brain areas to less well-studied behaviors can in the future be understood by measuring the way they format information from different sources.

We used artificial neural networks in a slightly different way than many recent neuroscience studies (*2, 45, 49–56*), because we used them simply as hypothesis generators. The use of RNNs has generated some controversy in neuroscience, mainly out of concern over the fact that the architecture of these networks is vastly different from biological networks (*57*). Our work highlights the value of using recurrent neural network models that are constrained by neural data to generate experimentally testable predictions about the relationships between neurons and behavior rather than as models of a biophysically realistic mechanism.

Finally, our study provides a resolution to the seeming paradox of having multiple brain areas that encode redundant sensory, cognitive, and motor information (‘everything is everywhere’). Our results suggest that different brain areas divide up neural computations in ways that are reflected in the formatting of different information sources. We demonstrate that the ability to decode information from a neural population does not imply much about how that population contributes to behavior. Instead, future efforts to understand neural computations and to intervene in disorders that affect cognition should focus on how the flexible reformatting of neural population responses in different areas enables flexible behavior.

## Supplementary figures

**Supplementary Figure 1:**
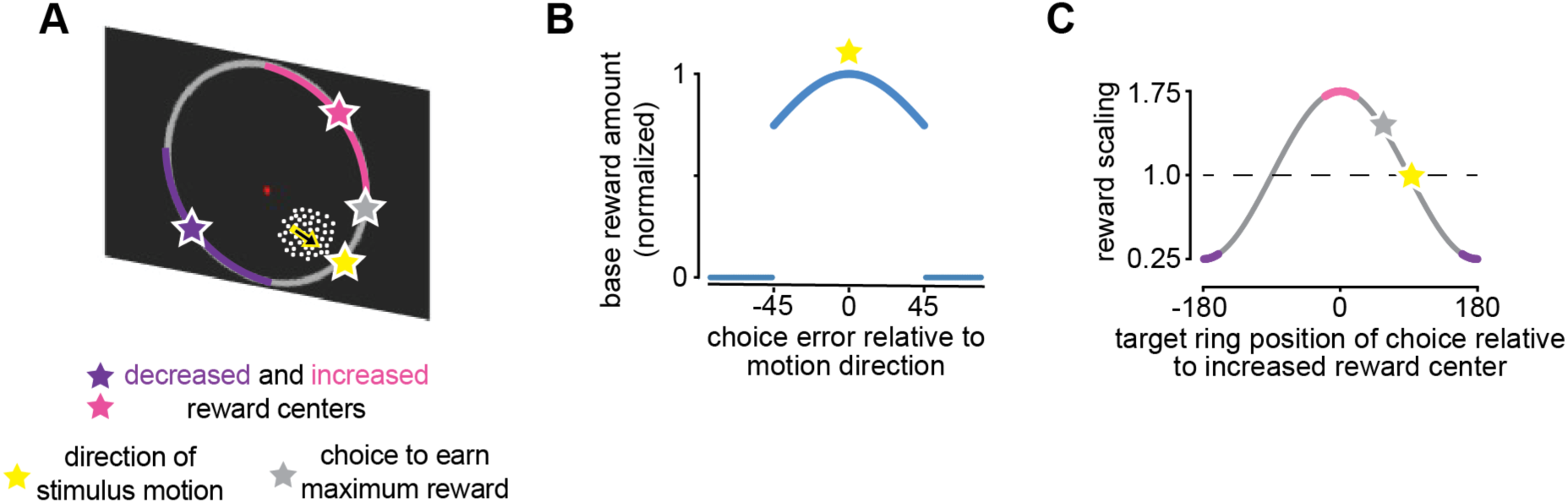
The monkeys were rewarded based on the accuracy of their motion direction judgment, increased or decreased by a reward scaling factor associated with different choices. A) Each trial required a motion direction judgment biased by reward scaling information indicated by the colored portions of the target ring. For the purposes of illustration, the reward scaling centers are noted with stars, the direction of stimulus motion is indicated with a yellow star and the choice associated with the largest reward is marked with a gray star. Both the motion direction and the reward condition were randomly interleaved from trial to trial, and the colors associated with increased or decreased reward varied in blocks with uncued changes. The raw reward amount varied across sessions, but rewards were always based on the (B) difference between the monkey’s choice and the direction of visual motion (with a hard cut off at +-45 degrees error) which was scaled by the values in (C). In this example reward condition, the increased reward center is the middle of the pink arc, and the decreased reward center is the middle of the purple arc.

**Supplementary Figure 2:**
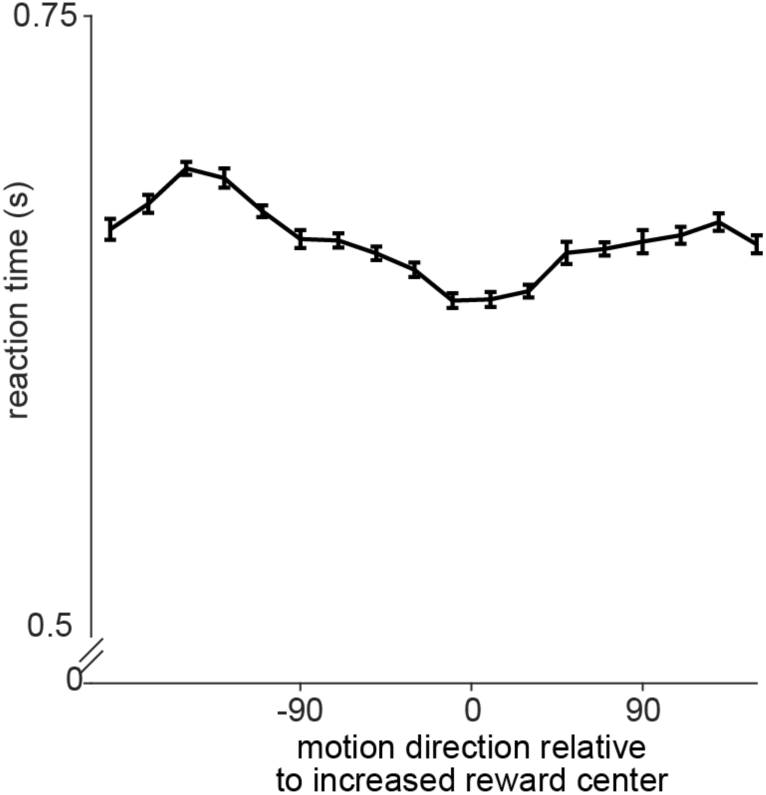
Reaction times during the continuous motion estimation task were slightly faster when the motion direction was aligned with the increased reward center. Mean reaction times (+-standard error of the mean) are plotted from 41,106 trials collected from 27 recording sessions (18 from monkey O). Reaction times are defined as the time from the onset of the visual stimulus to the time the monkey initiated the eye movement used to indicate a direction judgment. Reaction times were significantly modulated by the alignment between motion direction and the increased reward center (ANOVA, p<.01).

**Supplementary Figure 3:**
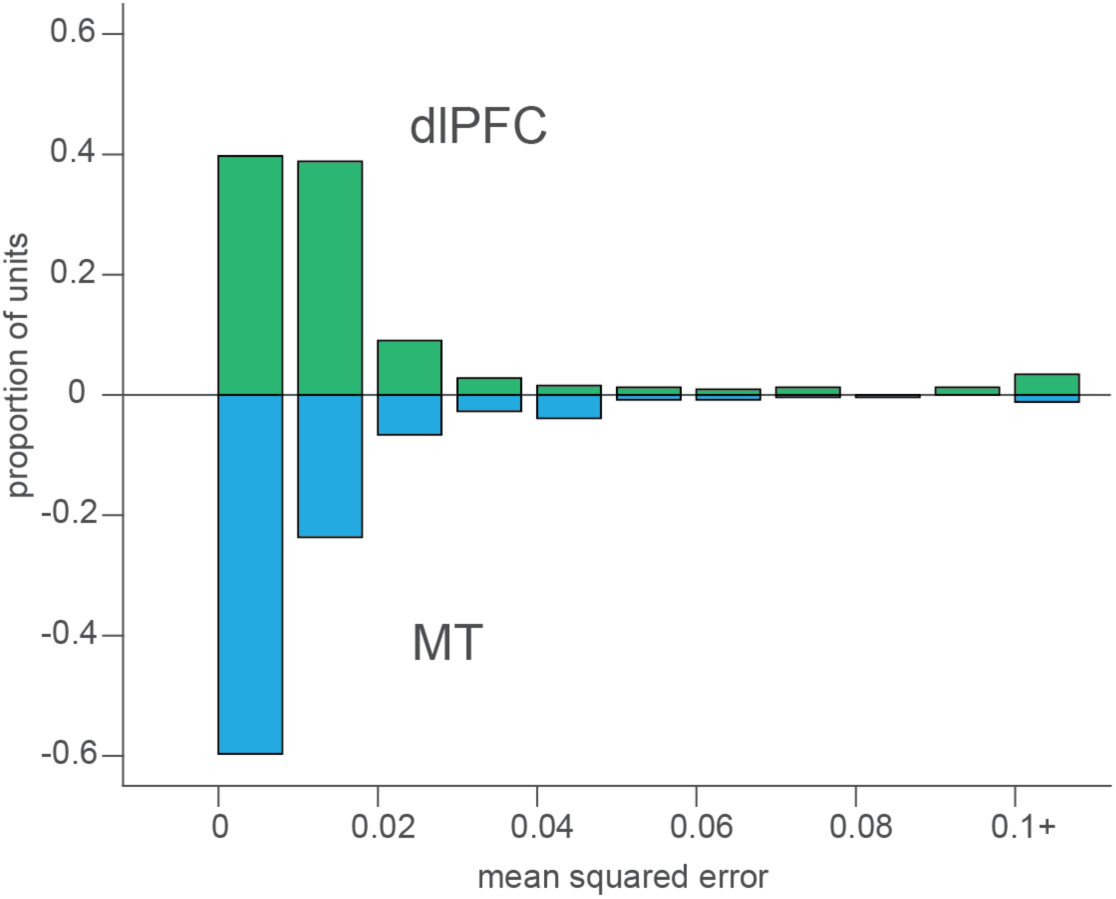
dlPFC units exhibit more non-linear mixed selectivity to visual motion direction and reward information than MT. The histograms of mean squared error of a motion direction tuning curve using a single scaling term to fit the two reward conditions (see Methods section 5.2) show greater error for dlPFC than MT (Wilcoxon rank sum test, p<.01 x 10^-8^). The histograms describe tuning curves for 258 MT units and 322 dlPFC units from 15 recording sessions in which there were simultaneous recordings in the two areas.

**Supplementary Figure 4:**
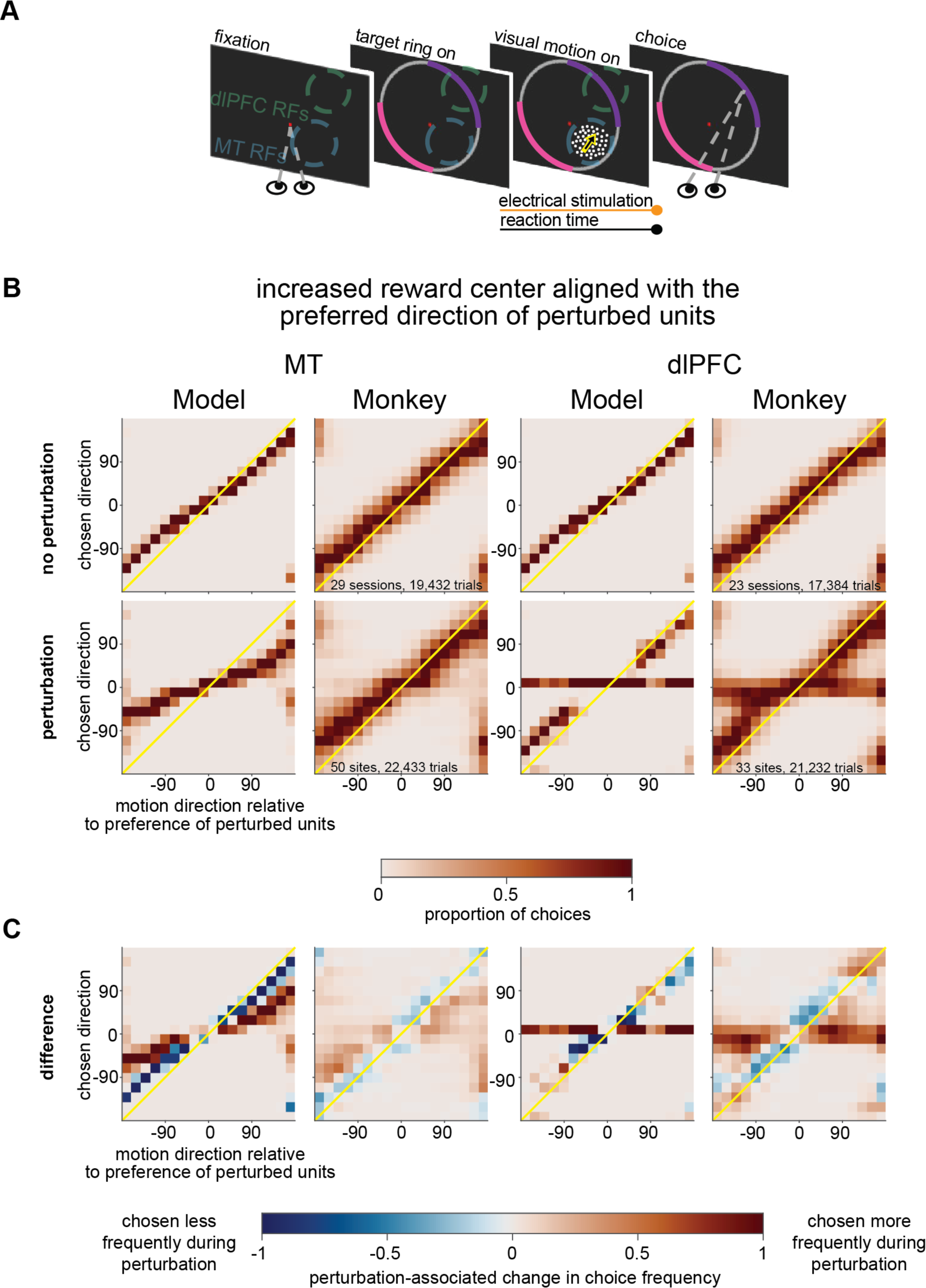
Behavioral choices from model and monkey during perturbation experiments when the increased reward center was aligned with the preference of the perturbed units. **A)** During monkey experiments, electrical stimulation coincided with the presentation of the visual stimulus. **B)** Summary of all choices by the model or monkeys when the increased reward center aligned with the preferred direction of the perturbed units. Data from 29 experimental sessions (comprising 50 sites; some sessions had stimulation on different electrodes on interleaved trials) of MT microstimulation (column 2), and 23 sessions (38 sites) of dlPFC microstimulation (column 4). Top row are choices during trials with no perturbation, bottom row are choices from trials with perturbation. All trial types, stimulus, and reward conditions were randomly interleaved within a session along with those shown in Supplementary Figure 5. Data are aligned to the tuning preference of the stimulated site, and sessions are only included in these analyses if the angular difference between that preference and the increased reward center was 30° or less. Data are normalized such that each column sums to 1. C) Plots reproduced from Figure 4B depicting the difference of the plots in (B).

**Supplementary Figure 5:**
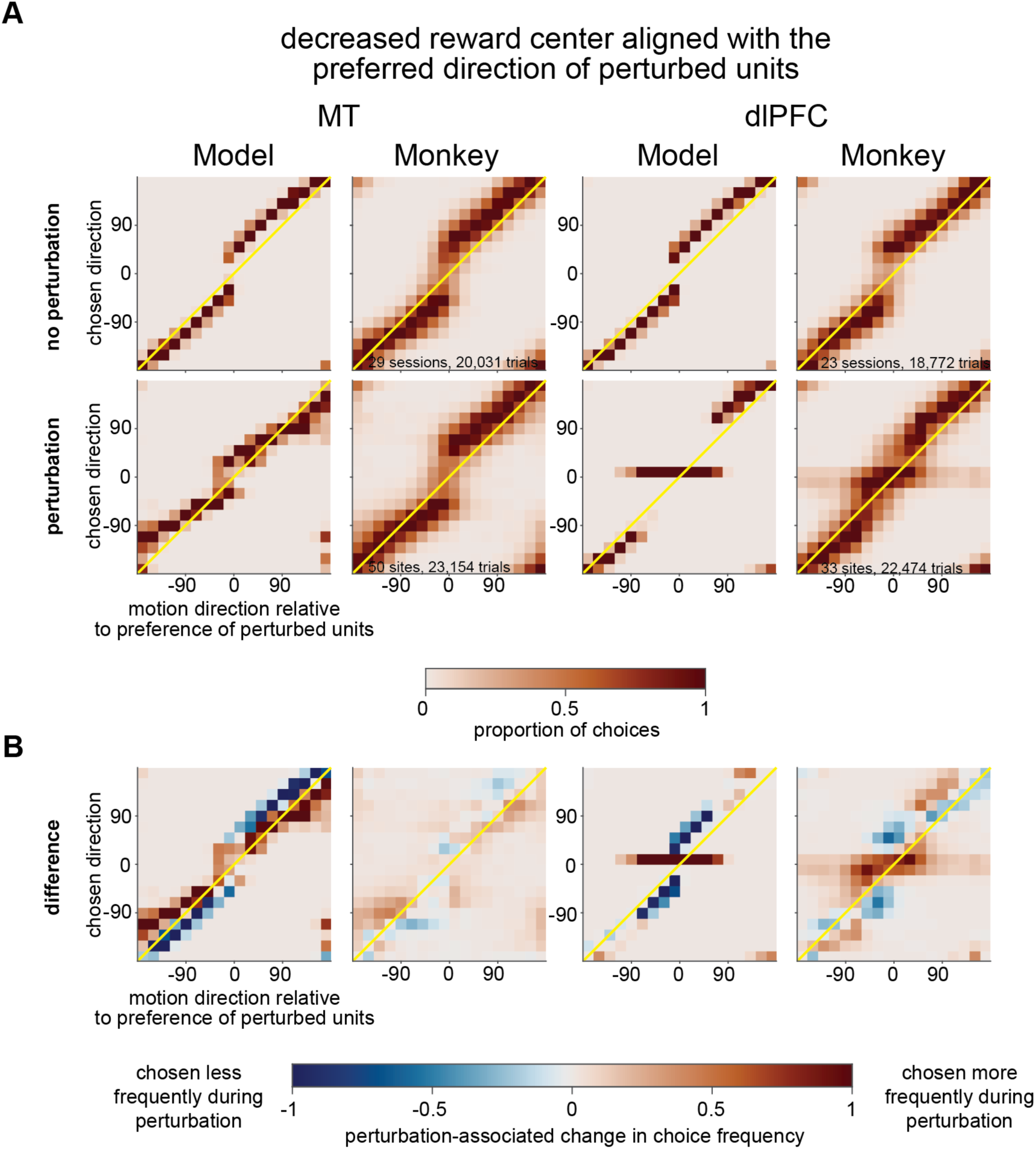
Behavioral choices from model and monkey during perturbation experiments when the decreased reward center was aligned with the preference of the perturbed units. **A)** Summary of all choices by the model or monkeys from the same experimental sessions and models as Supplementary Figure 4, on trials when the decreased reward center aligned with the preferred direction of the perturbed units. Conventions as in Supplementary Figure 4. B) Plots reproduced from Figure 4C depicting the difference of the plots in (A).

**Supplementary Figure 6:**
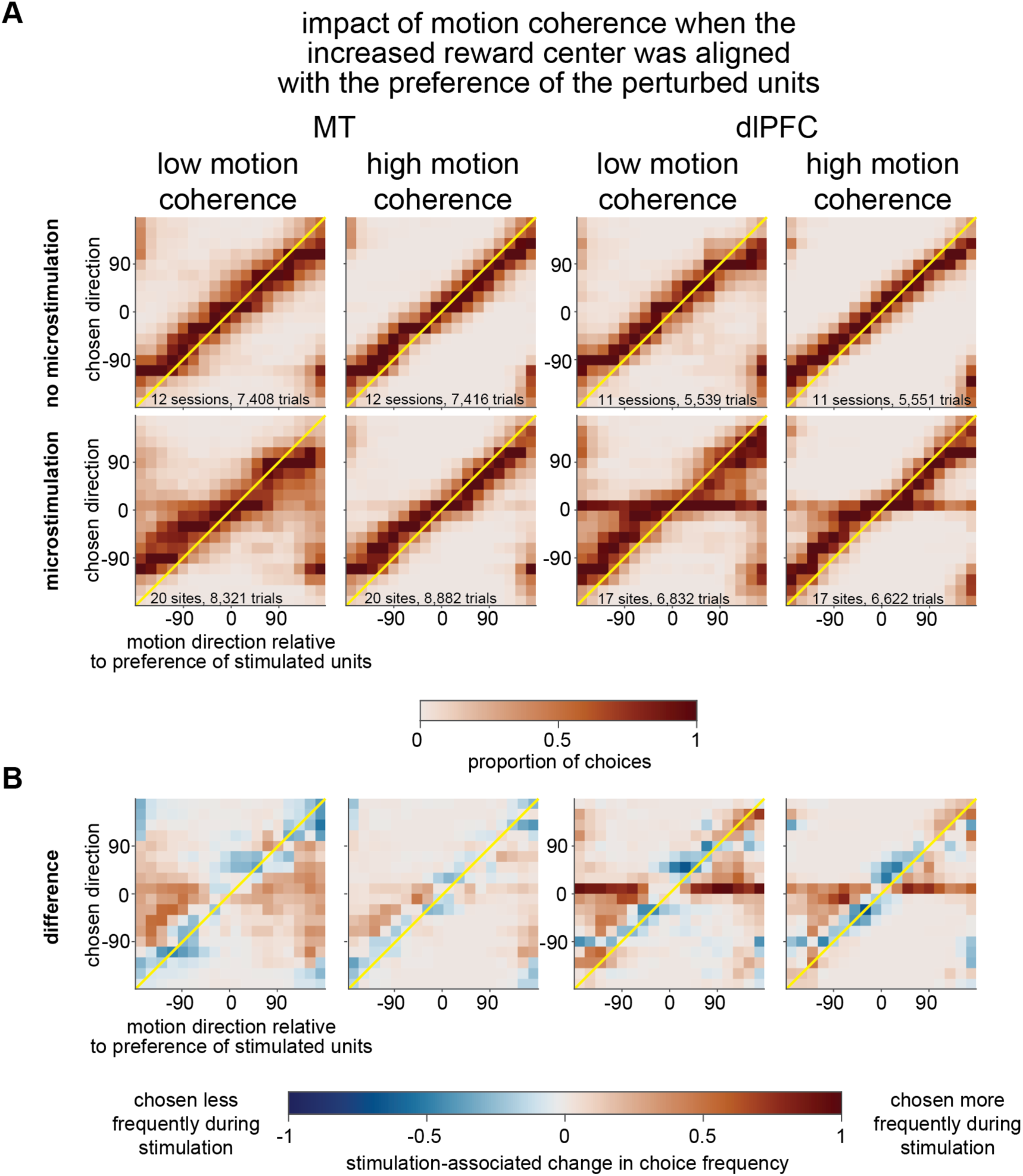
The magnitude of, but not qualitative difference between, the behavioral impact of MT and dlPFC microstimulation depends on motion strength, on trials when the increased reward center was aligned with the preference of the perturbed units. **A)** Summary of choices when the increased reward center aligned with the preferred directions of the microstimulated units during 12 experimental sessions (20 sites) of MT microstimulation and 11 sessions (17 sites) of dlPFC microstimulation using two motion coherences per session (low coherences were chosen between 15% and 20% coherence; high coherences were chosen between 35% and 50% coherence). The top row depicts trials with no microstimulation and the bottom row depicts trials with microstimulation. **B)** Microstimulation-related difference in choices in both areas. The proportion of choices from A during non-microstimulation trials were subtracted from the proportion of choices during microstimulation trials at each coherence level. In MT, when the visual motion stimulus was at a lower coherence, microstimulation biased choices more strongly toward the preference of the stimulated site. In dlPFC, when the visual motion stimulus was lower coherence, microstimulation elicited more choices near the preference of the stimulated site.

**Supplementary Figure 7:**
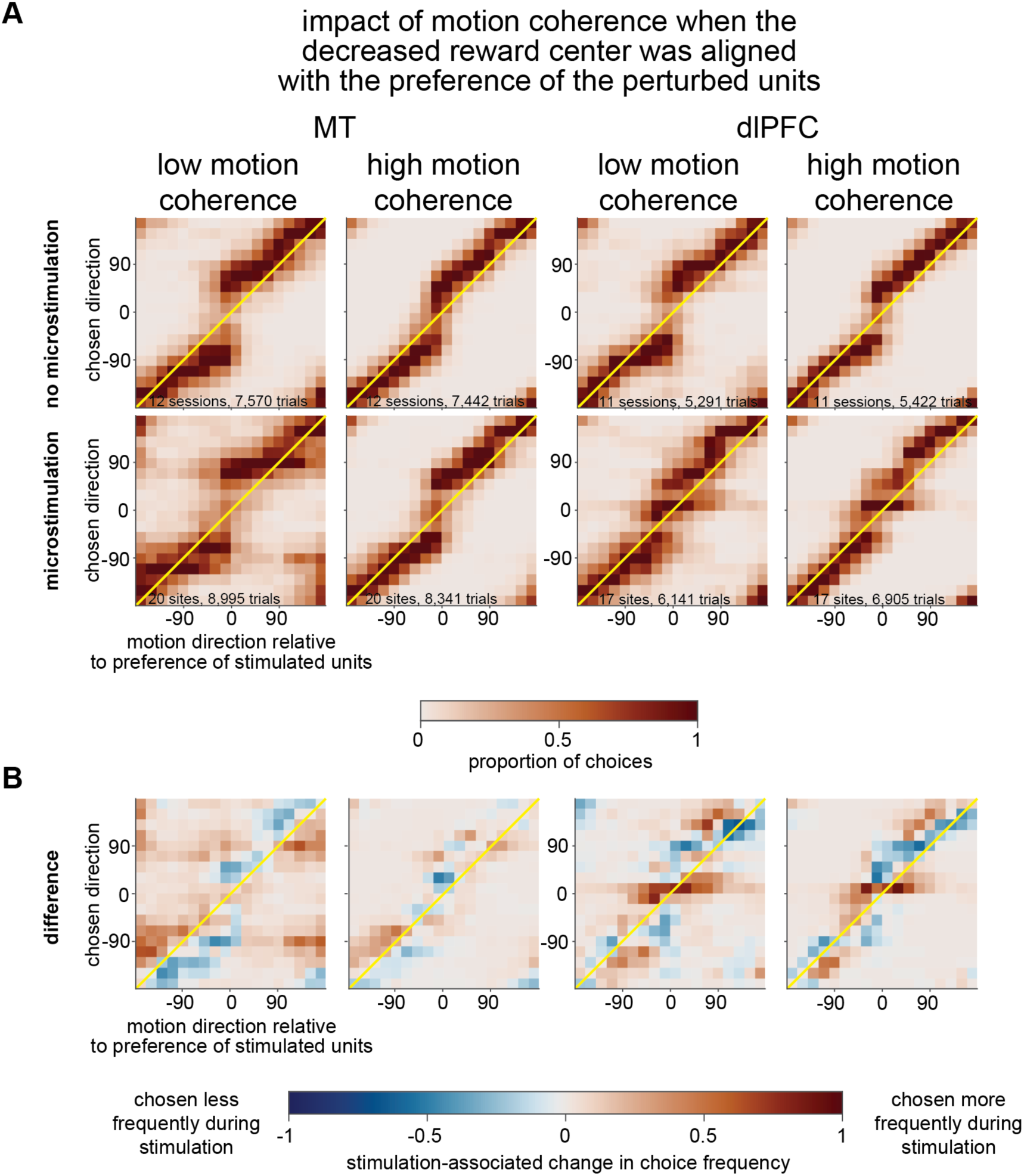
The magnitude of, but not qualitative difference between, the behavioral impact of MT and dlPFC microstimulation depends on motion strength, on trials when the increased reward center was aligned with the preference of the perturbed units. **A)** Summary of choices on trials from the same recording sessions and models as Supplementary Figure 6 when the decreased reward center aligned with the preferred directions of the microstimulated units. Conventions as in Supplementary Figure 6A. **B)** Microstimulation-related difference in choices in both areas. Conventions as in Supplementary Figure 6B.

**Supplementary Figure 8:**
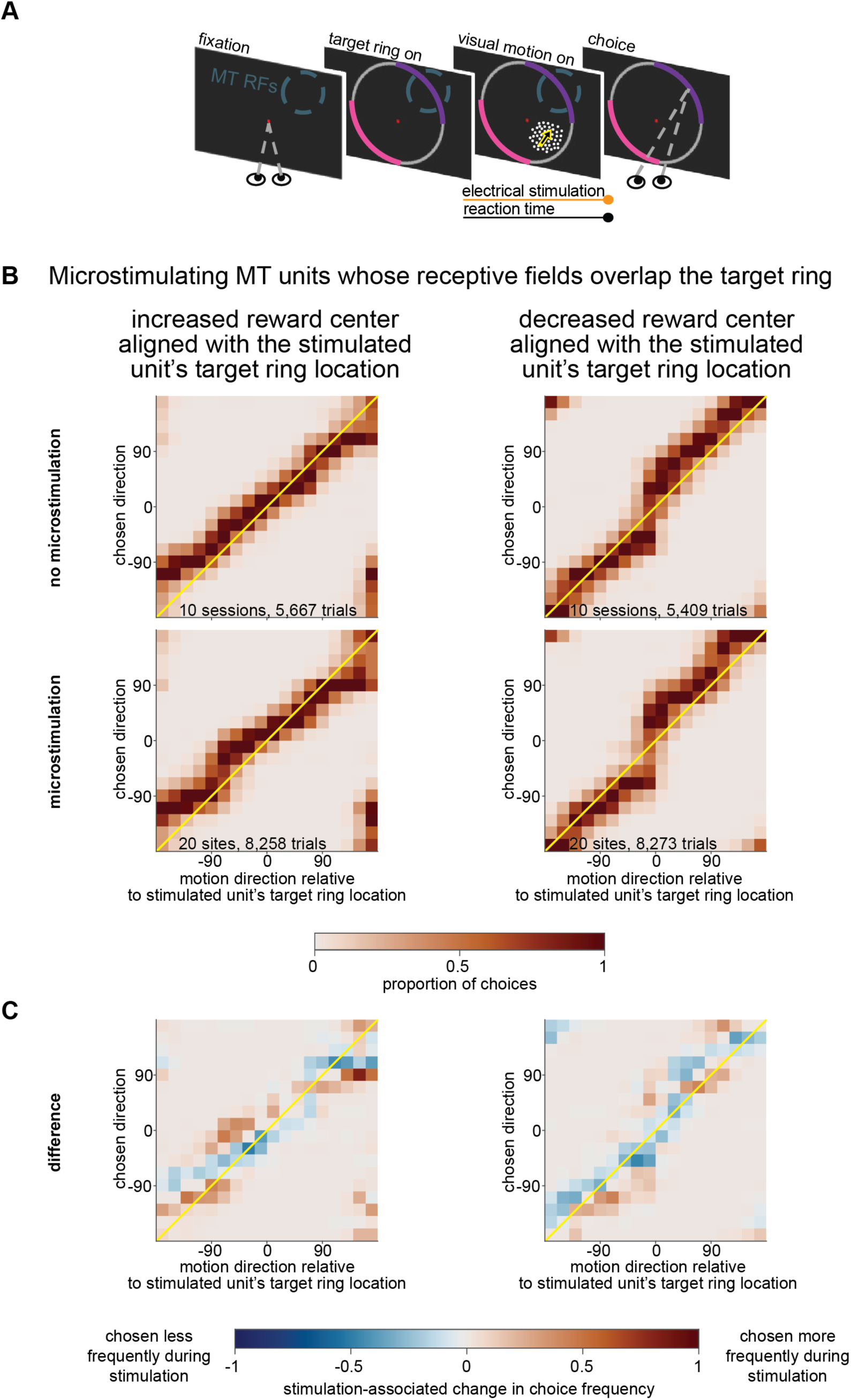
Microstimulation in MT does not bias choices when the target ring, not the visual motion stimulus, overlaps the receptive fields of the microstimulated units. **A)** As in all stimulation experiments, electrical stimulation coincided with the presentation of the visual stimulus, but unlike in most experiments (e.g. Figure 4), the receptive fields of the stimulated MT units overlapped the target ring, not the visual motion stimulus. **B)** Summary of choices from 10 experimental sessions (20 sites) of MT microstimulation when the receptive fields of the stimulated units overlapped the increased (left) or decreased (right) reward centers. Top row depicts trials with no microstimulation, bottom row depicts trials with microstimulation. **C)** Microstimulation-related difference in choices. The proportion of choices from B during non-microstimulation trials were subtracted from the proportion of choices during microstimulation trials for each reward condition. Choices were not biased by microstimulation in a way that is qualitatively similar to microstimulation in dlPFC (compare Figure 4B and C), even though in both cases, the receptive fields of the stimulated neurons overlapped part of the target ring.

## Acknowledgements

We are grateful to Karen McKracken for providing technical assistance, to Ramanujan Srinath, Benjamin Cowley and members of the Cohen lab for comments on earlier versions of this manuscript and other helpful suggestions. This work is supported by the Simons Foundation (Simons Collaboration on the Global Brain award 542961SPI) and the National Institutes of Health (awards R01EY022930, R01EY034723, and RF1NS121913).

## Author contributions

D.A.R. and M.R.C. designed the study. D.A.R. performed the experiments. S.M. and J.K. performed the recurrent network modelling. All authors analyzed data and wrote the paper.

## Competing interests

The authors declare no competing interests

## Notes

### Competing Interest Statement

The authors have declared no competing interest.

