## supplemental methods and figures for "Linking neural population formatting to function"

**Methods Contents**

1. Population formatting simulation
2. Subjects
3. Behavioral task
    3.1 Procedures
    3.2 Task and training
4. Electrophysiology experiments
    4.1 Procedures
    4.2 Recording locations
    4.3 Electrical microstimulation
5. Analysis of electrophysiological data
    5.1 Data inclusion criteria
    5.2 Mixed selectivity analysis
    5.3 Dimensionality reduction and angle calculations
6. Recurrent neural network model
    6.1 Training and architecture
    6.2 Model unit perturbation
7. Supplementary references

**1. Population formatting simulation**
We conducted a simple simulation to demonstrate that it is possible for populations of neurons with identical visual tuning functions and patterns of modulation by cognitive processes, like reward expectation, to format information from different sources in different ways. We simulated two populations of neurons, each of which consisted of units with the same motion direction tuning functions, which were Von Mises functions with uniformly distributed preferred directions (preferred directions varied in steps of 10°).

We simulated modulation by reward expectation in ways consistent with the small, heterogeneous, multiplicative gain changes observed in neural responses in tasks that vary reward expectation and feature attention (*1, 2*). We drew gains for each neuron from a uniform distribution between 1 and 1.2 and used these as a multiplier to the response of each neuron. Inspired by single neuron studies, we scaled this multiplier by the difference between the preferred direction of the neuron and the center of the high reward region. Depending on the extent to which these random gains happened to coordinate across the population, we could choose random draws such that the population formatted motion and reward information differently (Figure 1A-G).

**2. Subjects**
We trained two adult male rhesus monkeys, S and O (8 and 10 kg, respectively) to perform a continuous motion estimation task with variable reward expectation. Before training, both monkeys were prepared surgically with a custom titanium head holding device. To enable electrophysiological recordings, two recording cylinders (Crist instruments Co., Inc., Hagerstown, MD) were placed over 1) the principal sulcus and 2) visual cortex to provide access to dlPFC and MT, respectively. All surgical and behavioral procedures conformed to the guidelines established by the National Institutes of Health and were approved by the Institutional Animal Care and Use Committees of the University of Pittsburgh and Carnegie Mellon.

**3. Behavioral task**

**3.1. Procedures**
During training and recording sessions, the monkeys sat in a primate chair that provided head restraint. Visual stimuli were presented at a 120Hz refresh rate on a high-quality LCD monitor (VPixx Technologies Inc., QC Canada) placed 57cm from the monkey's eyes. Eye movements were monitored using an infrared optical eye tracker (Eyelink 1000, SR Research, ON, Canada). Behavioral control and stimulus presentation were managed by computers using a Unix based operating system (running MATLAB, The MathWorks, Inc., Natick, MA, and PsychToolbox (*3, 4*)).

**3.2. Task and training**

A trial began when the monkey fixated a central spot on a black background (inside an invisible 1° diameter window). Then a circular target ring was displayed and remained visible for the remainder of the trial. The target ring was uniform gray except for two segments 180° apart that were colored either pink or purple to signify the increased or decreased reward centers (see below and in Supplementary Figure 1). After a 200-400 ms delay, a dynamic random dot kinematogram was displayed in a stationary circular aperture inside the target ring. These motion stimuli were identical to those described in (*5*) and were presented at a single motion strength (coherence), typically between 40 and 50%, for most experiments (exceptions are described in Supplementary Figures 6 and 7). During multi-coherence experiments, trials using two motion coherences were randomly interleaved (low coherence: 15% - 20%; high coherence: 35% - 50%).

Monkeys were trained to make a saccadic eye movement to a location on the target ring to signify their choice. The monkeys could initiate the eye movement between 150 and 1500 ms after visual stimulus onset. The reward was based on the accuracy of the motion estimate, scaled by the amount indicated by the location of the pink- and purple- colored portions of the target ring. Which color signified the increased or decreased reward center switched in blocks with randomized, unsignaled changes throughout the experiment (typically 2-8 changes per session; Supplementary Figure 1).

During training, the monkeys first learned to perform the motion continuous estimation task before the reward scaling was introduced. The monkeys learned to saccade to a location on the target ring that matched the direction of motion of the visual stimulus, not where the vector of motion of the stimulus would cross the target ring. This design ensured that the monkeys' judgements would not be biased by the exact location of either the visual motion stimulus or the target ring, both of which could vary based on the receptive field locations of the recorded neurons.

### 4. Electrophysiology experiments

#### 4.1. Procedures

Neurophysiological recordings were performed using 24 or 32 channel linear probes (V- and S- probes; Plexon Inc, Dallas, TX) positioned using grids (Crist Instruments Company Inc., Hagerstown, MD) and advanced using a hydraulic microdrive (Kopf instruments, Tujunga, CA). Spiking activity, local field potentials, eye position, and task events were recorded at 30,000 samples/second (using Trellis software and Ripple recording hardware; Ripple, Salt Lake City, UT). All data were analyzed using custom scripts (MATLAB; The MathWorks, Inc., Natick, MA).

#### 4.2. Recording locations

Recordings in MT were made via a chamber positioned over visual cortex aimed anteriorly toward the middle temporal lobe. This posterior approach provided access to groups of MT units with similar spatial receptive fields but different direction tuning preferences across the contacts of a multi-channel probe (*6, 7*). MT units were identified based on stereotactic coordinates, gray-white matter transitions, and functional properties. In most experiments, we aimed for MT units with receptive fields that were less than 12 degrees eccentric so we could place the motion stimulus within their receptive fields without overlapping the target ring. When we electrically stimulated MT neurons whose receptive fields overlapped the target ring (Supplementary Figure 8), we used neurons with more eccentric receptive fields. During all recordings, we advanced the probe until we could stably record from as many visually responsive and direction-tuned units as possible. We measured spatial receptive field and direction tuning to verify electrode positioning, guide motion stimulus positioning, and, when relevant, identify the tuning preferences of sites recorded on the contacts we used for electrical stimulation.

Recordings in dlPFC were made via a chamber positioned over frontal cortex, anterior to the arcuate sulcus and spanning both sides of the principal sulcus (determined by stereotactic location and visual inspection during surgery). We recorded units along the banks and gyri of the principal sulcus. We used recording locations in which we observed spatial selectivity during any period of a delayed saccade task, and we stopped advancing the probe when we could stably record from as many selective units as possible. We typically recorded from units with response fields at similar eccentricities, whose locations sometimes progressed counter-clockwise as penetration depth increased. We preferred units whose response fields were more than 10 degrees eccentric so we could position a portion of the target ring within them and avoid overlap with the visual motion stimulus. We placed one colored portion of the ring at a location that was well covered by the response fields of the recorded units, and the other colored region was placed 180° away.

#### 4.3. Electrical microstimulation

During electrical microstimulation experiments, we selected two contacts on a multi-contact probe to be stimulated on randomly interleaved trials. Each trial consisted either of no microstimulation or stimulation on one electrode, and there was no statistical relationship between stimulation and the motion stimulus or reward condition. Microstimulation experiments were only ever performed in one area per session. Stimulation electrodes were randomly selected from suitable candidates based on tuning curves measured at the start of an experiment. MT sites were required to have a suitable visual receptive field location and strong direction tuning. dlPFC sites were required to have suitably positioned visual or premotor response fields.

Electrical microstimulation began with the onset of the visual motion stimulus and remained on until the monkey moved its eyes out of the fixation window to initiate a saccade. Microstimulation was biphasic and delivered at 200 Hz with an amplitude that ranged across sessions between 20 and 40 μA (*8-10*).

### 5. Analysis of electrophysiological data

#### 5.1. Inclusion criteria

MT units were included for analysis if their average firing rate during a 250 ms epoch following visual motion onset, lagged 50 ms for response latency, was at least 10% higher than during a period of stable fixation on a blank screen. dlPFC units were included for analysis if their firing rate during either of the two 250 ms epochs that followed the onset of the target ring or the visual motion stimulus, lagged by 50 ms, or the 250 ms that preceded eye movements was at least 10% higher than during a period of stable fixation on a blank screen. Trials were included for analysis if the monkey indicated a choice by looking at the target ring between 150 ms and 1500 ms after visual motion onset.

#### 5.2. Mixed selectivity analysis

We analyzed non-linear mixed selectivity for recordings from 15 sessions in which we recorded MT and dlPFC simultaneously. We evaluated the extent to which a linear scaling of motion direction tuning curves from one reward condition could explain the tuning curve in the other reward condition. We normalized the two vectors of mean responses from one unit from each reward condition such that their combined mean equaled 1. Next, we fit a line that scaled the vector from one condition to responses from the other and we evaluated the mean squared error in Supplementary Figure 3.

#### 5.3. Dimensionality reduction and angle calculations

To visualize the structure of population activity in both monkey (Figure 2) and model (Figure 3), we performed principal components analysis on trial averaged responses that were either averaged across motion direction conditions or reward condition. To determine the extent to which information about the two task features was represented in the same dimensions of neural population activity, we calculated the angle between the first principal components for motion direction and reward (Figure 2C).

### 6. Recurrent neural network model

#### 6.1 Training and architecture

The full neural network model consisted of two recurrent modules connected in series, such that part of the output of the MT-like module (module 1 in the equations below) determined the input to the dlPFC-like module (module 2). The modules were trained sequentially.

The dynamics of the modules' recurrent states, $x_1$ and $x_2$, are governed by the following equation:

$$x_{i,t+1} = \alpha\left(-x_{i,t} + A_i r_{i,t} + B_i u_{i,t} + b_{rec\,i}\right) + \sigma\sqrt{2\alpha}\, w_{i,t} \quad i = 1, 2$$

where the firing rate $r_{i,t} = \max\left(0,\ x_{i,t}\right)$ is the recurrent state passed through the rectified linear unit (ReLU) activation function, $w_{i,t} \sim N(0, 1)$ is a Gaussian noise process scaled by the recurrent noise factor $\sigma$, and $\alpha = \frac{dt}{\tau}$ is the ratio of the step size to the neural time constant. $A_i, B_i, b_{rec\,i}$ are the recurrent weight matrices, input weight matrices, and recurrent bias vectors of module $i$, respectively, and $u_{i,t}$ is the input to module $i$ at time $t$. The outputs of the modules are given by:

$$z_{1,t} = C_1 x_{1,t} + b_{out1}$$

$$z_{2,t} = C_2 r_{2,t} + b_{out2}$$

where $C_i, b_{out\,i}$ are the output weights matrix and output bias vector, respectively. The behavioral choice is taken to be $argmax(z_{2,T})$ where $T$ is the last time point of the trial.

For each timestep while the ring is present but before the onset of the motion stimulus, both the MT-like and the dlPFC-like modules received a two-dimensional input proportional to $[\sin\theta_R, \cos\theta_R]$ where $\theta_R$ is the increased reward center for that reward condition. The MT-like module outputs this same information, while the dlPFC-like module outputs a nonlinear function of these and its other inputs, as described below. Gaussian noise was added independently to all inputs at each timepoint, and no effort was made to match the variability in neural responses or in behavior to the experimental data. For each timestep during the motion stimulus, the MT-like module received a 72-dimensional motion input given by the von Mises function:

$$M_1(\theta_i, \theta_M) = -0.3 + e^{c*k_1*\cos(\theta_i-\theta_M)}$$

where $c \in [0, 1]$ and $\theta_M$ are the coherence and direction of the motion stimulus, and the 72 dimensions correspond to evenly spaced directions $\theta_i \in [0, 2\pi]$. The model in Figures 3 and 4 was trained with $c = 0.6,\ k_1 = 0.3$. The MT-like module was first trained separately to take in this noisy signal and output an amplified, sharper 72-dimensional motion output given by von Mises function:

$$M_2(\theta_i, \theta_M) = a * \left(-0.3 + e^{c*k_2*\cos(\theta_i-\theta_M)}\right)$$

It was also trained to output $[\sin\theta_R, \cos\theta_R]$ to ensure that the reward condition input is not simply ignored. After the MT-like module was trained, we froze the weights and trained the dlPFC-like module to take as inputs the motion output from the MT-like module and combine it with a reward condition input to produce a 72-dimensional reward function output given by:

$$R(\theta_i, \theta_M, \theta_R) = \begin{cases} \left(3.5 + 1.5 * \cos(1.8 * d_{\theta_i,\theta_M})\right) * \left(1 + 0.75 * \cos(d_{\theta_i,\theta_R})\right), & d_{\theta_i,\theta_M} \le \frac{\pi}{4} \\ 0, & d_{\theta_i,\theta_M} > \frac{\pi}{4} \end{cases}$$

where $d_{\theta_1,\theta_2}$ is the angular distance between $\theta_1$ and $\theta_2$. This is the same reward scaling as was used in the experiments (Supplementary Figure 1).

Both modules were trained by supervised learning with mean squared error loss using the Adam optimizer algorithm (*11*). The weights matrices were subject to a Dale's law constraint in which all input and output projections are excitatory (≥0), and 20% (30 out of 150) of recurrent units send solely inhibitory (≤0) projections while the remaining 80% send solely excitatory projections.

The training and testing of the neural network were implemented with custom code based on open-source packages (PsychRNN (*12*) and TensorFlow (*13*)).

#### 6.2 Model unit perturbation

To generate predictions about the impact of differences between the information representation in the MT-like and dlPFC-like module, we performed perturbation experiments in the fully trained models. We perturbed the activity of individual units in the recurrent layers of each module by adding an additional untrained input (that projected only to the stimulated unit) during the motion input period.
